# Modulating Intra-Nuclear LC3 with Small Molecules Rescues Cells from a Docetaxel-Induced Phenotype

**DOI:** 10.1101/2020.10.28.355826

**Authors:** Daniel P. Rosenberg, Likhitha Kolla, David S. Heo, Emily E. Cassio, Matthew J. Veenstra, Marianna Vakaki, Jibo Zhang, Cyril Anyetei-Anum, Lizabeth A. Allison, William J. Buchser

**Author notes:** <u>William J Buchser</u> (Corresponding), Department of Genetics, Washington University School of Medicine, 4515 McKinley Ave, St Louis, MO USA.

## Abstract

Nucleus-associated autophagy has been described as a cellular metabolic response by which nuclear material is actively degraded. This degradation occurs after stress, such as nuclear damage or the onset of tumorigenesis. Here we describe a nucleus-associated autophagic process distinct from other forms of selective autophagy in human cell lines. We found that although nuclear localization of MAP1LC3B (LC3) is not dependent on particular nuclear importins, knockdown of nuclear importins, which causes nuclear stress, can induce a nuclear autophagic response. Our characterization of this autophagic phenomenon was facilitated by chemical modulation of the process via two compounds discovered previously in a high content analysis. These small molecules bidirectionally regulate nuclear LC3 in human renal, pancreatic, and bladder cell lines. One molecule (NSC31762 or *DTEP*) *enhances* nuclear LC3 puncta and increases lysosomal targeting of LC3. This compound also decreases the nuclear envelope protein LaminB1. Another molecule (NSC279895 or *DIHI*) *reduces* the nuclear localization of LC3. Finally, we applied these chemical tools in the setting of mitotic-disruptor induced nuclear stress. The compound DIHI, shown to reduce nuclear autophagic puncta, diminished the mitotic disruptor effect. These new tools will allow for deeper exploration of nucleus-associated autophagies, and could serve as proof-of-principle in guiding new therapies for diseases involving nuclear stress.

## Introduction

Autophagy is the systematic recycling and destruction of cellular components such as ineffectual organelles and is a basal process in most eukaryotes^1^. Selective forms of autophagy have been described as highly specific, under clear regulation, and have been found to span cellular compartments such as the mitochondria (mitophagy), the endoplasmic reticulum (ER-phagy), and the peroxisome (pexophagy)^2–4^. Nucleophagy, or degradation of nuclear material via an autophagic pathway, has been identified as a unique process in yeast and mammalian systems^5^. Recent studies have confirmed the existence of nucleophagy and have correlated it with diverse cellular processes including epidermal differentiation^6^, senescence after DNA damage^7^ and the clearing of extra-nuclear DNA^8,9^. However, the set of conditions under which nucleophagy is activated, the identification of the degraded nuclear substrates, and the mechanism of transport are still under investigation.

Nucleophagy appears to be a stress response system in line with the basic functions of other autophagy processes. It is recognized that the nucleus has complex pathways in place which respond to stress or damage^10–12^. Normally, proteins enter and exit the nucleus mediated by transporters that are members of the importin/exportin family^13^. Disruption of nuclear transport through the nuclear pore complex leads to nuclear stress and subsequent disease phenotypes, including age-related damage, cancer, and neurodegeneration^14–17^. Here we investigate whether nucleus-associated autophagy is upregulated in response to nuclear-transport stress, and how nucleocytoplasmic shuttling of protein products might be affected.

The loss or malfunction of autophagy can lead to cellular damage and the onset of many diseases including neurodegeneration and cancer^18^. It has been shown that oncogenic mutations in HRas lead to mis-regulated cell division and can trigger nucleophagy, leading to the removal of the nuclear envelope and structural proteins such as nuclear LaminB1. The end result of this nucleophagic process is the induction of cellular senescence and prevention of further damage to the rest of the tissue^7,19^.

When DNA damage occurs in tandem with malfunction or loss of nuclear autophagy, the risk of tumorigenesis and cell death increases^20^. The remarkable ability of cancer cells to withstand nuclear stress as a consequence of abnormal ploidy^21^ raises questions as to whether these cells manipulate nuclear forms of autophagy to prolong survival. Here, we look at a more extreme case as proof-of-principle by using mitotic-inhibitor induced polyploidy^22,23^ (MDIP) to model nuclear stress. We seek to better understand the interplay between nuclear autophagy and nuclear stress secondary to MDIP.

Here, we utilize two new chemical modulators of LC3B localization^24^ to better understand nucleus-associated autophagy. Given that nuclear autophagic puncta are infrequently observed in untreated cultured cells, these modulators up- or down-regulated expression of puncta allowing for tangible characterization of the process. Our characterization shows that the autophagic protein Microtubule Associated Protein 1 Light Chain 3 Beta (MAP1LC3B, hereafter referred to as LC3) localizes to regions of the nucleus with low DNA content (nuclear holes), and on occasion with large amounts of extra-nuclear DNA (micronuclei). Furthermore, our results suggest nuclear LC3 localization is increased under nuclear stress induced by knockdown of nuclear import proteins. Finally, we use the chemical modulators to investigate the functional role of this form of autophagy and to mitigate nuclear stress in the setting of DNA stress caused by microtubule inhibitors.

## Methods

### Cell Culture and Transfection

Human cancer cell lines from ATCC (Manassas, VA) were used: renal 786-0 (CRL1932), CAKI-1 (HTB46), RCC4 (CVCL0498), pancreatic cell line PANC-1 (CRL1469), colon cancer HCT 116 (CCL-247), and human bladder cancer cell line T-24 (HTB-4). Cells were cultured in DMEM (Thermofisher 11995-065) supplemented with FBS (Thermofisher 26140079) and PenStrep (Thermofisher 15140-122) in a humidified incubator at 37°C with 5% CO2. Cells were seeded in 96-well plates at 3,000 cells per well in 100μL of supplemented media. Plates were incubated at 37°C for 24 hours prior to manipulation.

Pre-designed SureSilencing™ short hairpin RNA (shRNA) plasmids from SABioscience (Frederick, MD) were used in order to target specific mRNA of importin β1 (KPNB1), importin 7 (IPO7), importin 8 (IPO8), and a scrambled negative control^25^. For time lapse experiments, we used the RFP-GFP-LC3B plasmid^26^ (gift from Earl Godfrey) and RFP-GFP-LaminB 1 (gift from Zhixun Dou). eGFP-LC3 was also used (Addgene 111546^27^), packaged in lentivirus using PSPax2 and VSVg in HEK293 cells. At 70% confluency, we transfected cells with shRNA plasmids or RFP-GFP-[] plasmids using Lipofectamine 2000 (Thermofisher 11668-027) and the manufacturer’s protocol. Ten hours after transfection, media was replaced and the cells were treated with the chemical compounds. After three hours, the compounds were rinsed out with fresh media, and the cells were incubated for 24 hours at 37°C.

### Nucleus assays

Heterokaryon nucleocytoplasmic shuttling assays were performed as described^25,28–30^. In brief, HeLa (human) cells were transfected with GFP-TRα1 then fused with untransfected NIH/3T3 (mouse) cells using 50% polyethylene glycol 1500. Fused cells were incubated for 4 hours at 37°C with cycloheximide (to prevent de novo protein synthesis) and either DMSO (vehicle control), DIHI, or DTEP. The human nucleus was distinguished from the mouse nucleus by differential coloration with Hoechst. To visualize heterokaryons, cells were fixed and stained for actin with Vectashield containing TRITC-phalloidin (red). Shuttling was viewed by epifluorescence microscopy.

For the polyploidy experiments, cells were treated with 0.1ug/mL colcemid (Thermofisher 15210040) for three hours followed by a hypotonic shock with 75mM KCL, and fixed with 3:1 ratio of methanol:glacial acetic acid fixative. A few drops of the fixed cell suspension were placed onto microscope slides and air-dried. The slides were then stained with Giemsa G-banding stain and rinsed in Giemsa buffer (Thermofisher 10092013, 10582013). Confocal microscopy was used to capture 1μm ‘sections’ of nuclei. Fields were then manually scored for cells in metaphase and anaphase. Nuclear material in these phases were then manually measured to count chromosomes and identify the presence of polyploidy.

### Immunofluorescence

Following treatment and incubation, cells were fixed with 3.2% paraformaldahyde (ThermoFisher) in PBS. Plates were blocked and permeabilized in one step using bovine serum albumin (BSA) and 0.1% Triton X-100 in PBS at room temperature for 1 hour (both from Sigma-Aldrich). Antibodies were diluted in a blocking mixture consisting of a 1:1 ratio of PBS and BSA. Nuclear material was stained with Hoechst (ThermoFisher, H3570) at a 1:5000 dilution and endogenous LC3 was tagged with an LC3B/MAP1LC3B primary antibody (ThermoFisher, L10382) used at a dilution of 1:500 for 72 hours at 4°C. Other antibodies use include the LaminB1 (ThermoFisher, PA5-19468) 1:650, and LAMP2 Antibody (ThermoFisher, H4B4 MA1-205) 1:200. After incubation, the primary antibody was rinsed three times with PBS then Alexa Fluor 546-tagged goat anti-rabbit IgG secondary antibody (ThermoFisher, A-11035) was added at a 1:1250 dilution and incubated for 1 hour before a final 3x rinse with PBS.

### Microscopy

Endpoint experiments were visualized using a Nikon inverted epifluorescence microscope. A 40x DIC objective was used with the DAPI (400-800 msec exposure) and TRITC (800-1600 msec exposure) filters. Microscopy was semi-automated such that four different images were acquired in each well (using *multi-point)* and the focal plane for each well was determined by eye.

For confocal microscopy, cells were cultured on glass coverslips (#1.5), treated for a period of 4 hours, and subsequently fixed and stained after 24 hours. Coverslips were inverted and mounted onto glass slides using Fluoromount (Sigma-Aldrich). A Nikon confocal microscope was used to acquire Z-stacks at an average of 8-10 steps over a range of 6-9μM (top to bottom of the nucleus). The pinhole was set to 1.7 A.U., while laser power was held constant across samples.

For time-lapse experiments, 786-0 cells were transfected with an RFP-GFP-LC3 plasmid. After 8 hours, DMEM was replaced with FluoroBright DMEM (Thermofisher, A1896701) and cells were treated with autophagy-modulating compounds. Temperature, CO2, and humidity were kept constant in a live cell chamber. Images were captured via confocal microscopy at 20minute intervals over 16 hours using a pinhole of 1.2 A.U. and 4x line averaging.

### Image Analysis

Images were processed in *FIJI Imaged*^3^. *ImageJ* was used to create maximum intensity projections of confocal z-stacks, z-slice views (dynamic reslice), and plot intensity profiles. Large 2×2 tiled images gathered from the inverted microscope were split into four by a script running in *ImageJ.* Open-source software *CellProfiler* was used for automated image analysis^32^. The *CellProfiler* pipeline identified individual nuclei via Mixture of Gaussians (MoG) thresholding, which then allowed for analysis of intensity, localization, and prevalence of LC3 and nuclear material within the cell. The image-based flat files returned from *CellProfiler* were compiled, and *Spotfire DecisionSite* (Tibco Spotfire) was used to “quality control” the tracing by examining each of the traced wells for errors. The scripts for *ImageJ* and *CellProfiler* are included in the **Supplement**. Images that contained fewer than 3 cells per image, contained an error in tracing, or contained other artifacts were omitted from the overall analysis. When replicates from different plates (often from different cultures and days) were analyzed, they were first normalized so that the global mean of all the wells for all parameters studied were equal to 1 (by dividing by the global mean for each plate). To analyze the correlation between the two channels in cells, the Coloc2 program in *ImageJ/FIJI* was used on two slices selected from a z-stack of confocal images.

### Statistics

Statistics were performed on at least three biological replicates, defined as coming from different culture flasks. Overall alpha was set to 0.05. For experiments with multiple conditions, ANOVA was performed first, and then post-tests were run if overall ANOVA was significant. All analysis of dose effects used linear regression.

### Protein Analysis

Knockdown of importin β1, importin 7, and importin 8, compared to scrambled control shRNA, and protein compartment analysis was done by western blot as previously described^25^. Briefly, 786-0 cells in 6-well plates were transfected with sets of four shRNA plasmids (SABioscience, Frederick, MD) and lysed 24 hours post-transfection. Antibodies were used with the following concentrations: anti-GAPDH (Santa Cruz Biotechnology Inc, Dallas, TX), 1:5000; anti-importin β1 (Santa Cruz), 1:2000; anti-importin 7, 1:1000; anti-importin 8, 1:250 (both Abcam, Cambridge, MA). Independent replicates of each importin knockdown were completed from different cultures across different days.

## Results

### Compounds Manipulate Nuclear LC3 Puncta

Functional experiments from a previous study^33^ analyzed chemical compounds for nuclear LC3 phenotypes. Two distinct phenotypes arose after treatment with two of these compounds. The first molecule increased nuclear LC3 intensity; NSC31762, 2,6-**D**iiodo-4-[2-(1,3,3-**T**rimethylindol-2-ylidene) **E**thenyl] **P**henol (abbreviated ***DTEP***). Nuclear LC3 in cells exposed to DTEP at increasing concentrations (**Fig 1a**) had a significantly positive regression between DTEP concentration and LC3 accumulation within the nucleus (Linear regression, p =1.5×10^−5^, F=58). DTEP is itself fluorescent at 590 nm; while this did not present any issues in our assays, it raises the possibility for further study using this molecule as a novel nuclear-associated autophagy tracer. The second compound decreased nuclear LC3 intensity; NSC279895, 1-(2,4-**Dihy**droxyphenyl) Heptane-1,2-Dione (abbreviated ***DIHI***). There was a significant decrease in nuclear LC3 intensity as DIHI concentration increased (**Fig 1b,** Linear regression p=0.0081, F=7.8).

**Figure 1.**
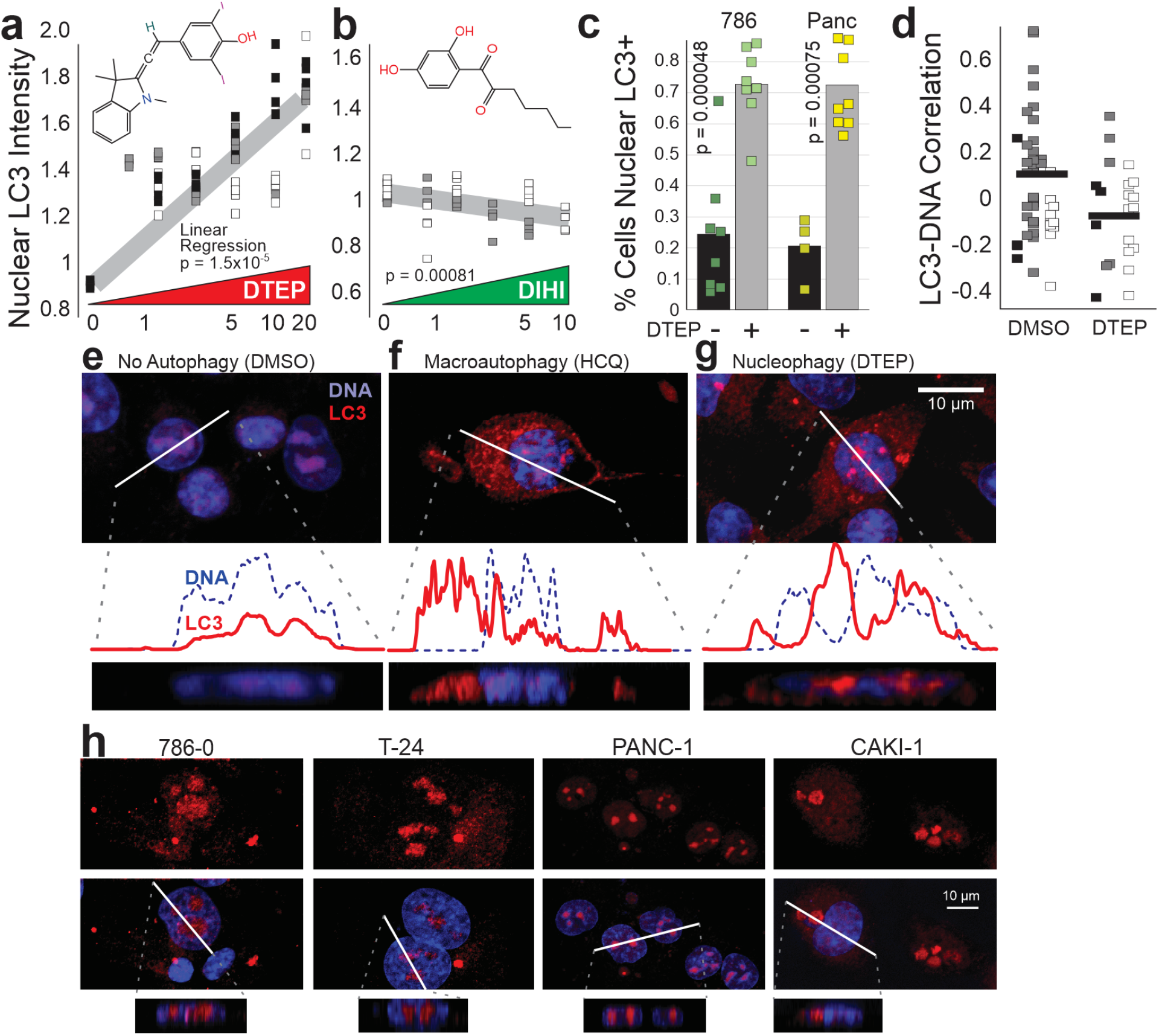
Compounds that Enhance and Inhibit LC3 Nuclear Puncta. **a**) Increasing concentrations of the compound DTEP (structure pictured) in 786-0 cells resulted in a significant increase in nuclear LC3 compared to a DMSO control (Linear regression, p=1.5×10^−5^). **b**) Increasing concentrations of the compound DIHI (structure pictured) in 786-0 cells results in a significant decrease in nuclear LC3 intensity (p=0.0081). Markers represent well average, shadings indicate different biological replicates. **c**) Markers show % of cells above background levels of nuclear autophagic puncta (4 biological replicates each). Bars demonstrate the average % cells per condition. The presence of DTEP significantly increases the percentage of cells that are positive for nuclear LC3 between both pancreatic and renal cell lines (p < 0.001, t-test of biologically independent replicates). **d**) Nuclear LC3 puncta have variable amounts of colocalization with nuclear DNA (p = 0.048, dots represent the LC3-DNA pixel-pixel based Pearson’s correlation of individual cells analyzed from 3 different cell lines; white: RCC4, gray: 786-0, black: CAKI-1). e-g) Upper panels are confocal slices on 786-0 cells. A line plot (generated based on the solid white line in the upper panels) indicates topographical association between DNA (blue) and LC3 (red). Below the line plot, a scale-matched depth reconstruction (z-stack) is shown. **e**) 786-0 cells treated with DMSO (vehicle) show low autophagic activity and weak topographical association. **f**) Cells treated with hydroxychloroquine (HCQ) have classic perinuclear localization of LC3. **g**) Treatment with the compound DTEP results in higher concentrations of LC3 inside the nucleus. Line scans show the anti-correlated nature of the DNA and LC3 (with LC3 localized to low-DNA nuclear holes). **h**) Confocal microscopy showing nucleophagy across renal clear cell carcinomas (786-0, CAKI-1), bladder cancer (T-24), and pancreatic carcinoma (PANC-1). Z-stacks show LC3 (red) accumulated within the nucleus (blue).

Previous examination of the autophagic signature of a number of cancer cell lines revealed that renal and pancreatic cells in particular were more likely to express nuclear LC3 puncta under both normal and stress conditions, and thus were used in this experiment^34^. While a fraction of control cells possess nuclear autophagic puncta, it is a relatively rare occurrence (**Fig 1c** black bars, **Fig S3**). LC3-associated micronuclei were also observed (**Fig S1**). Ultra-structurally, electron micrographs of nuclei appeared normal in the presence of DTEP and DIHI (**Fig S2)**. DTEP’s ability to induce nuclear puncta was analyzed in both 786-0 cells and PANC-1 cells over several biological replicates. DTEP increased the fraction of cells with nuclear LC3 by 1.5X (cells with average nuclear LC3 intensity above DMSO control levels, **Fig 1c** gray bars, t-test, p < 0.0001).

Presence of LC3 within the nucleus (not above or below) was confirmed using multi-step z-stack confocal microscopy. To determine whether the localization of the LC3 within the nucleus was consistent in the DTEP condition, a colocalization analysis was performed. Twenty-five representative images from biological replicates across three cell lines were analyzed. Individual nuclei were analyzed by confocal slice, and the correlation between LC3 and DNA was slightly lower in DTEP-treated cells than control (**Fig 1d**; t-test, p = 0.048). The variation between cells, however, was high, indicating that colocalization between these nuclear holes and LC3 puncta may depend on additional factors.

We next asked how this chemical-induced phenotype compared to other autophagy-related phenotypes. 786-0 human renal cells were exposed to diluent (DMSO), macro-autophagy was blocked using hydroxychloroquine (HCQ), and nuclear puncta were invoked using DTEP (as above). Confocal microscopy confirmed little to no LC3 in the cytoplasm or in the nucleus of most control cells (**Fig 1e**). When treated with HCQ, nonspecific extra nuclear macroautophagy was clear (**Fig 1f**). There was little to no association between nuclear DNA intensity and LC3 intensity within the nucleus; most LC3 remains in the cytoplasm, as is seen both in the line scan and in the z-stack. When cells were treated with DTEP, prominent nuclear LC3 puncta appeared (**Fig 1g**). In the representative cell shown, LC3 & DTEP were localized to regions of low DNA density / nuclear holes.

We asked whether DTEP’s induction of nuclear punctae was conserved across cell types. Treatment with 20 μM DTEP produced a strong increase in nuclear LC3 in multiple cell lines, including renal lines 786-0, and CAKI-1, T-24 bladder cancer, as well as the pancreatic line PANC-1 (**Fig 1h**).

We also directly confirmed that exogenous LC3 could be influenced by DTEP using a GFP-LC3 virus (we evaluated several to find this construct with near-endogenous expression). DTEP was able to increase the fraction of LC3-GFP+ cells (**Fig S3a**) and we could also confirm a clear non-overlap between cells that had taken up large amounts of DTEP (which fluoresces at 590 nm) and cells with LC3-GFP (i.e. not a fluorescence overlap, **Fig S3b**). LC3 was also probed in a protein analysis after DTEP and DIHI treatment in both the cytoplasm and the nucleus (**Fig S4)**. DTEP was found to subtly increase LC3 compared to the DMSO control. This was inconsequential, however, given the small number of cells with this specific phenotype. We conclude that DTEP and DIHI represent new chemical tools to modulate LC3 localization.

### Nuclear Autophagic Puncta Represent a Physiologic Intra-Nuclear Process

To determine the significance of this DTEP-induced intra-nuclear LC3, we investigated the degradation of nuclear envelope protein LaminB 1 using time-lapse microscopy in the presence of DTEP and DIHI (**Fig 2a)**. To monitor nuclear flux, we first used the tandem RFP-GFP-LaminB1 reporter, in which acid sensitive GFP is quenched in a low pH environment, usually the lysosome, leaving only red fluorescence as an indicator of completed autophagic activity^26^. Both the DMSO control and DIHI treated cells showed only a moderate decrease in the normalized green fluorescence. In the presence of DTEP, we instead observed a sharp decrease in the amount of green fluorescence intensity after six hours, suggesting DTEP-induced degradation of the nuclear envelope protein. Z-stacks of the nucleus confirm the DTEP-induced decrease in fluorescence within the nucleus over this time course (**Fig 2b**). The rate of the green fluorescence intensity change (slope) was significant for the DTEP treatment group (**Fig 2c**, ANOVA p = 0.0005, DTEP p = 0.0003). The ratio of GFP/RFP was not significant, leading us to further experiments to determine the cause of the degradation. We asked whether LaminB1 could be found in the cytosol after DTEP treatment, in particular after the addition of Bafilomycin to inhibit lysosomal turn-over. We found the cytosol to be clouded with LaminB1 staining in the DTEP conditions with Bafilomycin, and we also carefully quantified the cytosolic localization and found significantly more LaminB1 staining in the presence of Bafilomycin and DTEP (**Fig S5**, ANOVA p = 0.0117). It is clear that both exogenous and endogenous LaminB1 is reduced from the nuclear lamina after DTEP treatment and is subsequently found in the cytosol and near LAMP2 puncta.

**Figure 2.**
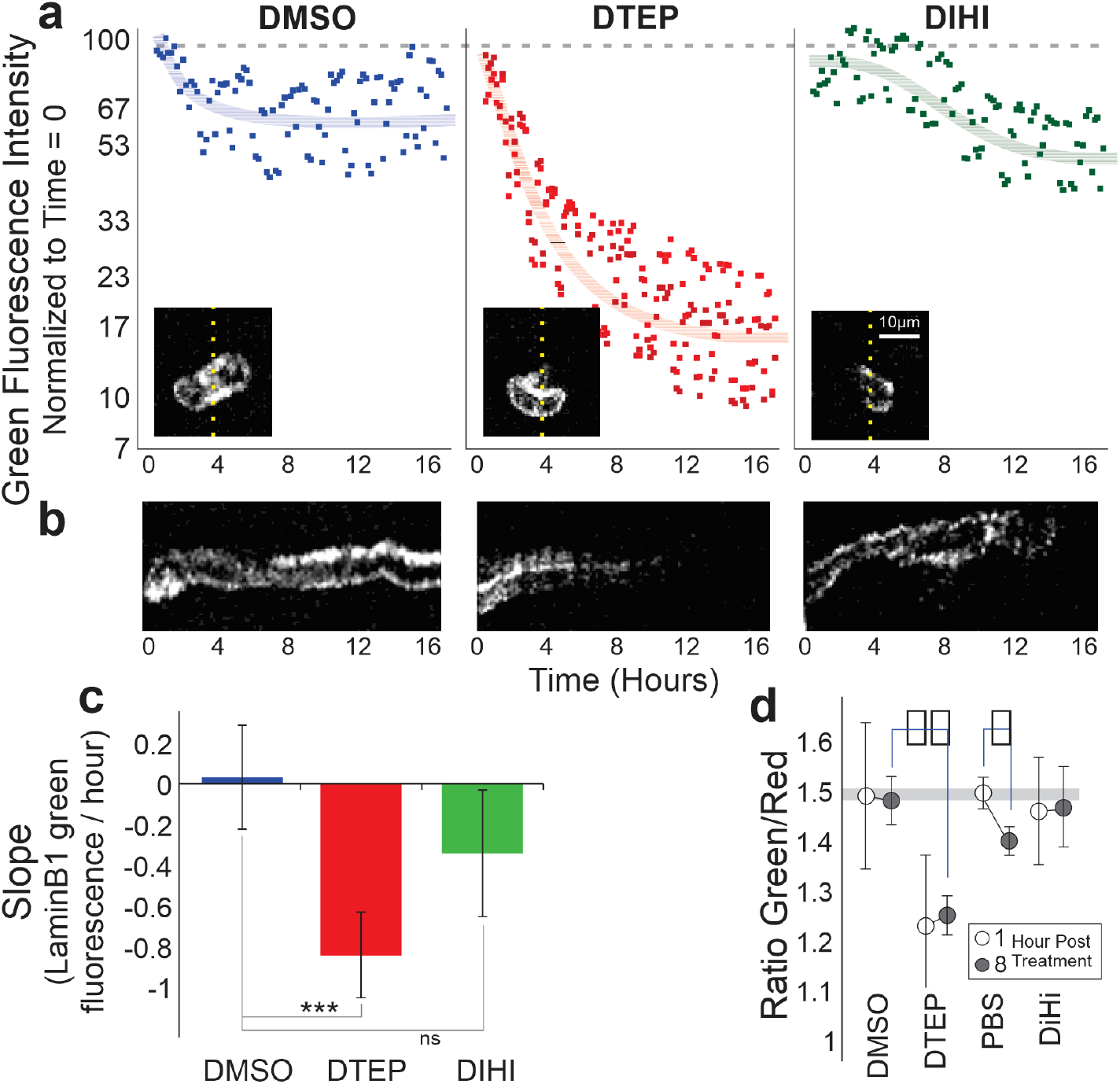
Degradation of LaminB1 and LC3 after Compound Treatment. **a**) Degradation of LaminB1 over time in the presence of DTEP and DIHI was monitored via confocal microscopy over the course of 16 hours. DMSO and DIHI did not result in a significant decrease in green fluorescence (decrease indicates the presence of the LaminB1 fusion protein in the lysosome), while DTEP caused a sharp drop around hour 4. **b**) Kymographs generated from the Z-stack slice indicated in (**a**) with the dotted line. These depict the change in green fluorescence over the course of 16 hours. **c**) Slope of LaminB1-GFP change over time (hours) (ANOVA p = 0.000502 F = 15.28, 9 cells/field, DMSO: 8 fields, DTEP: 4 fields, DIHI: 3 fields). **d**) 786-0 cells transfected with a tandem RFP-GFP-LC3 sensor were monitored for degradation over the course of eight hours. Markers show average ratio of green/red fluorescence with standard deviation between cells. Starvation (PBS) results in an increase in red signal (LC3 in the autolysosome, p = 0.037). Cells treated with 10 μM compound DTEP exhibit a significant increase in autophagy, especially eight hours after treatment (p = 0.00011).

In addition to looking for the degradation of the nuclear component (LaminB1), we also sought to quantify LC3 degradation. We ask whether DTEP accelerates the rate of autophagy with an RFP-GFP-LC3 reporter (GFP is quenched by lysosome). This reporter expresses LC3 at supra-physiologically levels, enhancing nuclear LC3 in most cells. We tested the degradation of this tagged LC3 as a proxy for autophagic rate. Treatment with vehicle (DMSO) or DIHI showed no significant change in GFP/RFP ratios over the course of an eight-hour period. A relative decrease in GFP, however, was observed in both the starvation treatment (PBS t-test p=0.037), as well as the 10μM DTEP treatment, even after 1 hour (t-test p=4.2×10^−6^) (**Fig 2d**).

### Loss of Importins Induce Nuclear LC3 Localization

We hypothesized that disrupting importin-dependent nucleocytoplasmic shuttling (nuclear transport) may induce nuclear stress and enhance nuclear autophagy^35^. We tested this hypothesis by knockdown of nuclear importins β1, 7 and 8. LC3 intensity within the nucleus was measured and compared to a scrambled shRNA control. Nuclear LC3 intensity significantly increased among importin β1, 7 and 8 knockdown cells compared with scrambled shRNA or untreated controls (**Fig 3a**, 2 wells each across 4 biologically independent replicates. ANOVA p=0.000141, f=6.91, post t-test p = 0.00045, p = 0.014, p = 0.016 respectively). The loss of importin7/8-mediated nuclear transport may have triggered nuclear stress and subsequent nucleus-associated autophagy. Knockdown efficiency was confirmed via western blot (**Fig 3b**).

**Figure 3.**
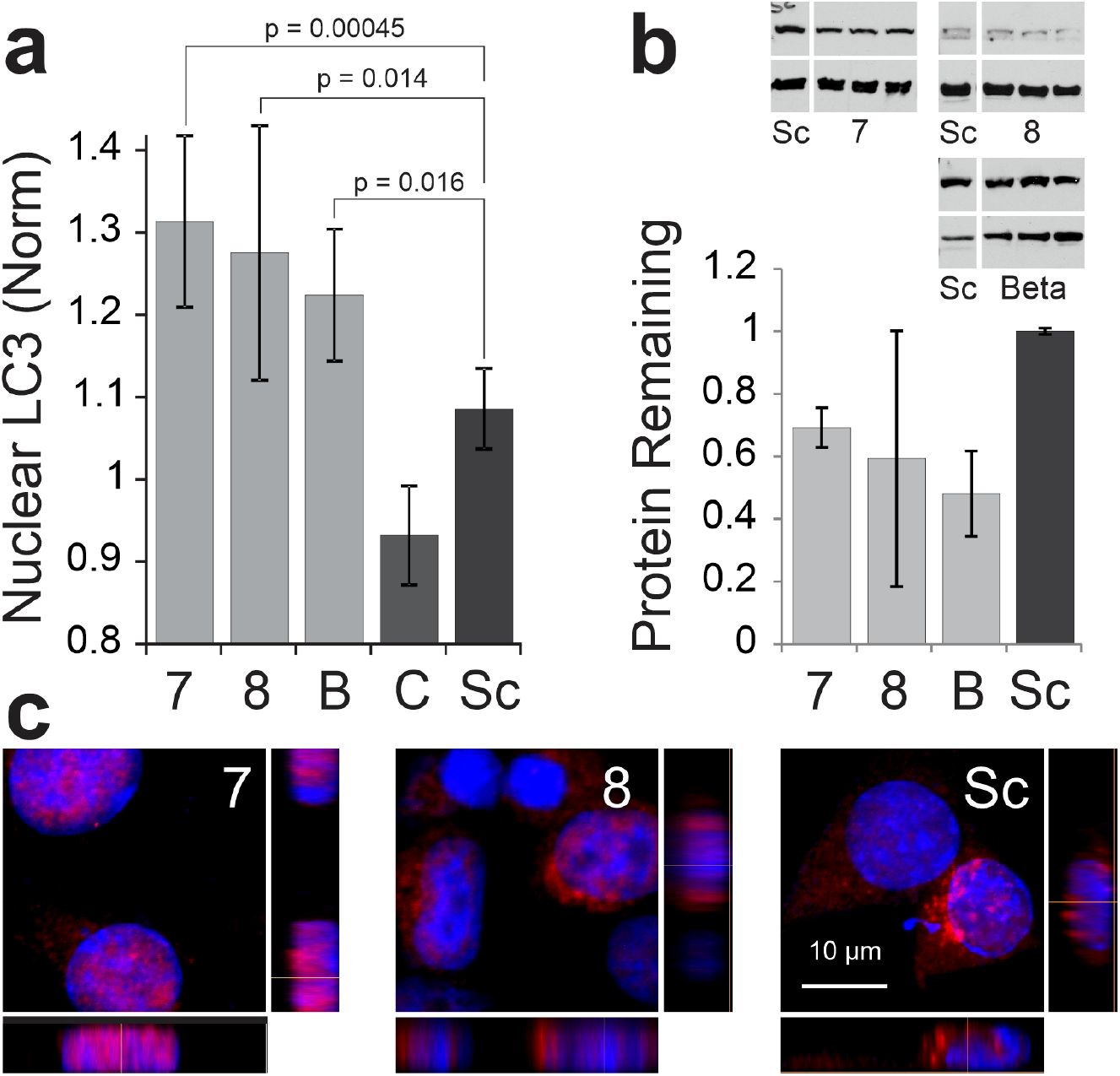
Knockdown of Several Importins Leads to Heightened Nuclear LC3 Accumulation. **a**) Average Nuclear LC3 intensity of 786-0 cells after knockdown of importins 7, 8 and β1, compared with untreated control and scrambled (Sc) shRNA. Error bars show standard deviation. **b**) Western blots (importins in upper bands and GAPDH in lower bands) and densitometric quantifications of knocked down importin 7, 8, and β1 when compared to a scrambled shRNA (average and standard deviation of 7 biological replicates). Western membranes are cropped as indicated by the white gaps and full membranes are shown in Fig S7. **c**) Micrographs show a maximum projection and two reconstructed side views of 786-0 renal cells transfected with shRNAs against importin 7, 8 or scrambled shRNAs. Cells were stained for LC3 (red) and DNA (blue). Importin 7 knockdown cells have the clearest intra-nuclear puncta (especially evident when viewed from the side), whereas importin 8 and Sc have both cytosolic and nuclear LC3 distribution.

Confocal microscopy was used to determine whether the localization of LC3 was juxta-nuclear or intra-nuclear after importin knockdown (**Fig 3c**). Importin 7 knockdown resulted in prominent intra-nuclear localization of LC3 whereas Importin 8 knockdown had both intra- and juxta-nuclear LC3.

Biochemical methods were used on cellular fractions to determine the effects of DTEP and DIHI on importin mediated transport of nuclear LC3. Treatment of cells with DTEP and DIHI resulted in virtually no change in importin expression in both nuclear and cytosolic fractions (**Fig S4**, no significant differences).

Finally, we wanted to find out if the compounds might directly manipulate transport across the nucleus, perhaps inducing autophagy through the same mechanism as importin7/8 knockdown. Additional protein analysis was used to quantify the levels of importins 8a, 8b, B1, as well as LC3a and LC3b (**Fig S4**). The quantity of protein within the nuclear fraction was nearly undetectable, indicating insignificant results regarding the effects of DTEP and DIHI on these nuclear transport proteins.

A heterokaryon assay was used to investigate the effects of our compounds on nucleocytoplasmic shuttling of protein products. We looked at the movement of thyroid hormone receptor α1 (TRα), which is known to rapidly shuttle between the cytoplasm and nucleus, as a marker of nuclear transport efficiency. Neither DTEP nor DIHI had any effect on nucleocytoplasmic shuttling compared to the DMSO control (**Fig S6).** Together, these results imply that this nuclear autophagy may respond to deficits in nuclear transport, however, the compounds have no direct effects on transport.

### Modulating Nuclear Autophagy Can Revert a Docetaxel-Induced Phenotype

The mitotic disruptors docetaxel and nocodazole halt cell division but lead to the abnormal phenotype of polyploidy and cellular senescence (which we will call mitotic-disruptor induced polyploidy or MDIP). Since it has already been shown that nucleophagy is important in the setting of senescence, we used MDIP as the paradigm in which to test the functional significance of DTEP and DIHI manipulation of nuclear autophagic puncta.

Quantification of the DNA content revealed that the majority of docetaxel or nocodazole-treated 786-0 renal cancer cells have nearly double the amount of DNA compared to DMSO-exposed control cells, indicating disrupted mitosis and polyploidy (**Fig S7, S8**). When 786-0 cells were exposed to different amounts of docetaxel (up to 3.2 μg/mL), the cell density decreased as the dose increased, indicating either cell death or reduced cell division (**Fig 4a**, linear regression, p=1.02×10^−5^, f=44.8). This decrease in cell density was complemented with an increase in average nuclear area (p=1.78×10^−5^). A second chemical, nocodazole, was used to induce mitotic disruption via G2/M arrest. Nocodazole treatment (up to 30 μg/mL) also lowered cell density and caused the nuclear area to expand (**Fig 4d**, linear regression density p=2.9×10^−5^, f=39.2, area p=0.0067, f=10.4), indicating reduced cell division due to mitotic blockade. As expected, mitotic disruptors produced a distinct nuclear phenotype (MDIP) and restricted cell division.

**Figure 4.**
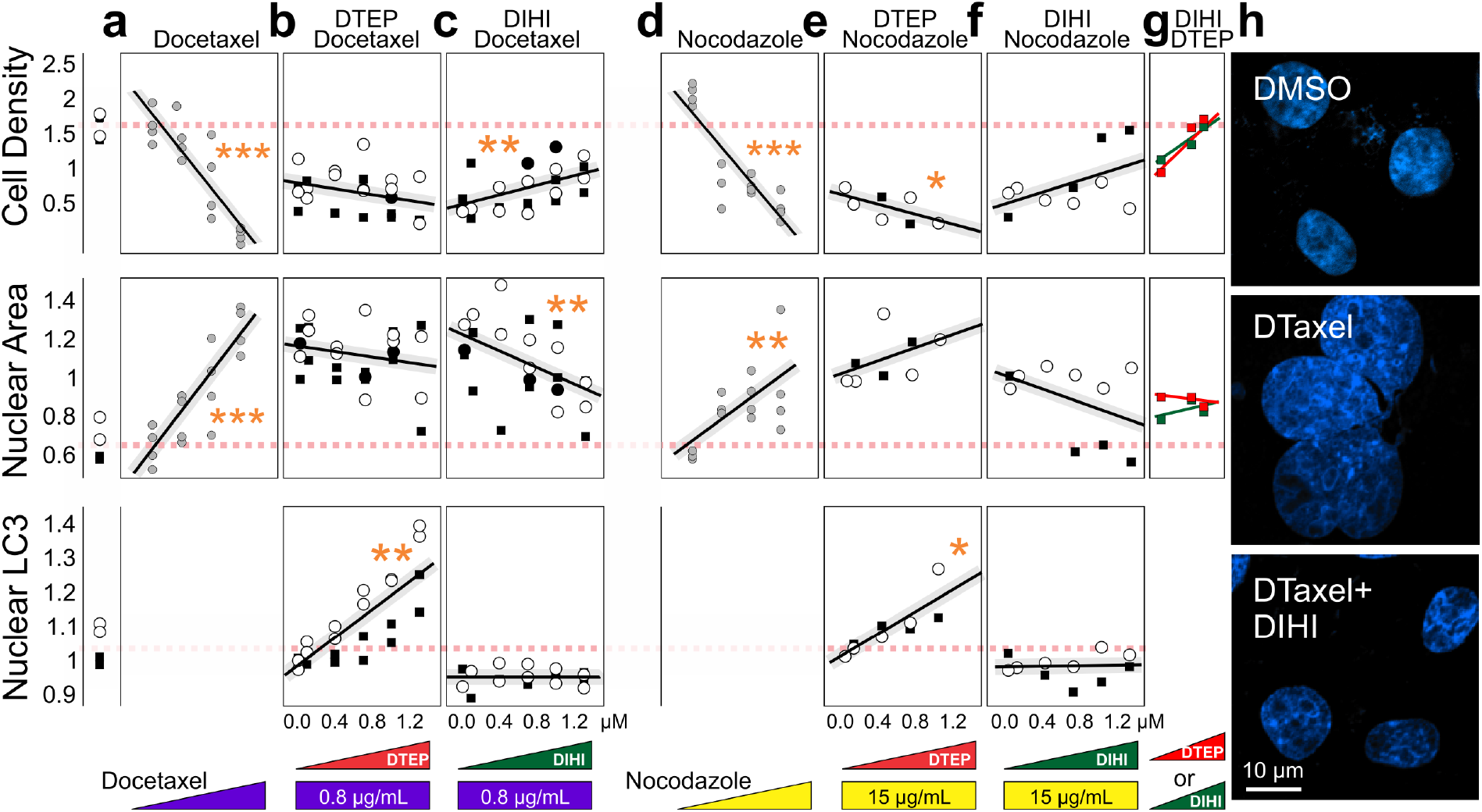
Compounds DTEP and DIHI effect Docetaxel- and Nocodazole-treated cells. **a-f)** 786-0 responses after exposure to various treatments. Upper panels show cell density, middle panels show nuclear area, and bottom panels show nuclear LC3 intensity. Markers represent normalized well averages from independent experiments (indicated by different marker types). Control levels of cell density, nuclear area, and nuclear LC3 are indicated by the red dashed lines traversing the panels. **a-c**) Cells treated with docetaxel, **d-f**) cells treated with nocodazole. **a)** Increasing docetaxel dose reduces cell density, and increases nuclear area increases (asterisks indicate significance in a linear regression). In the presence of background docetaxel, DTEP (**b**) or DIHI (**c**) were added in increasing doses. **d)** A nocodazole gradient lowers cell density and increases nuclear area through mitotic blockade. Increasing doses of the compounds DTEP (**e**) and DIHI (1). **g**) Gradients of the DTEP or DIHI alone (not significant). **h**) Representative confocal images of 786-0 renal cells exposed to DMSO (top), docetaxel 0.8 μg/mL (DT, middle), or docetaxel and 15 μM DIHI (bottom).

We next tested the LC3-localization modulators DTEP and DIHI in the setting of the mitotic disruptors. 786-0 renal cells were treated with 0.8 μg/mL of docetaxel for three hours and then exposed to an increasing DTEP concentration series (up to 1μM, **Fig 4b**). In the presence of docetaxel-induced polyploidy, an increased concentration of DTEP resulted in an increase in nuclear LC3 (bottom panel, p = 0.0047, f=16.7). DTEP was unable to interfere with the MDIP, and even contributed to further cell density reduction (**Fig 4b** upper panel). Conversely, 786-0 cells pre-treated with docetaxel and increasing concentrations of DIHI showed a significant increase in cell density (**Fig 4C** upper panel, p=0.0042, f=12.0) and decrease in nuclear area (middle panel, p=0.0043, f=11.9), indicating a possible reversion of MDIP. Representative images of nuclei after treatment with docetaxel and DIHI are shown in **Fig 4h** bottom panel.

We then asked whether DTEP or DIHI were able to interfere with the effects of nocodazole by exposing 786-0 cells to 15μg/mL nocodazole and a subsequent DTEP or DIHI treatment. DTEP concentration and nuclear LC3 were found to have a positive relationship as seen in previous experiments, but DTEP was unable to reverse nocodazole’s phenotypic effects (**Fig 4e)**. When DIHI was added in the presence of 15 μg/mL of nocodazole, there was no *significant* change in LC3, area, or cell density (**Fig 4f**). Neither DTEP nor DIHI alone had significant effects on the nuclear area or the cell density (**Fig 4g**).

To better document the chromosomes and DNA content in the setting of MDIP, cells were arrested at metaphase and anaphase with 0.1ug/mL colcemid (**Fig S8)**. In the absence of docetaxel, DIHI has no effect on the chromosome number or DNA content compared to the DMSO control, but docetaxel causes a large increase in the DNA content of many cells. Subsequent addition of DIHI significantly reduces the number of cells with this phenotype. Furthermore, DMSO and DIHI treated cells show a negligible percentage of polyploid cells. On the contrary, many of the docetaxel-treated cells were polyploid, but the percentage of polyploid cells was significantly reduced when docetaxel treatment was followed by DIHI treatment. We conclude that DIHI, which we showed to inhibit nuclear autophagic puncta, restores the cells from docetaxel-induced cell loss and nuclear expansion.

## Discussion

While some studies have documented nuclear autophagy as a cellular phenomenon, none have been able to specifically manipulate the process. We have shown here that nuclear-LC3 localization is a process phenotypically distinct from macroautophagy, likely related to nucleophagy and may be implicated in the survival of cells under nuclear stress. Additionally, we have utilized two recently-identified small molecules (DTEP, DIHI) to regulate the amount of LC3 that is present within the nucleus. We attempted to elucidate the relationship between nuclear stress and nuclear-LC3 localization by knocking down components of the nuclear import machinery. As expected from previous studies^35^, nuclear importins 7, 8, and β1 do not appear to play a role in LC3 nuclear entry. Instead, the knockdown of these importins led to nuclear stress which induced nuclear autophagic puncta. The presence of nuclear LC3 across multiple cell lines suggests that the process is not an artifact confined to a specific subset of cells^36^, rather, a distinct stress response pathway that is likely conserved. Nuclear LC3 is associated with an increase in LC3 found in nuclear holes. In the setting of MDIP, we found that treatment with DIHI, a potential inhibitor of the nuclear LC3, results in an increase in cell numbers and decrease in DNA content, reversing the effects of docetaxel.

Nucleus-associated autophagy likely encompasses a collection of varied cellular mechanisms with diverse functions that may span the activation of senescence, disposal of micronuclei, and perhaps even removal of incongruous strands of DNA^7^. Our ability to control and disrupt specific forms of nuclear autophagy is critical to understand their function and mechanism, while simultaneously leveraging the process for therapeutic applications. We hypothesized, in accordance with Dou *et al,* that upregulating nucleophagy may cause an increase in lamin-degradation, resulting in the high occurrence of nuclear autophagic puncta observed. We have discovered an application of a chemical tool, DIHI, which may also inhibit this lamin-degrading nucleophagy, thus preserving the nucleus, allowing for continued cell division. Our data confirms modulating autophagy (via DTEP) effects the turnover of LaminB1 (Figure 2), but the pathway which transports nuclear LC3 to the nuclear envelope remains unknown. It is possible that the tandem RFP-GFP-LC3 (or LaminB1) used to measure autophagic activity could encounter acidic environments outside of the lysosome that would cause degradation and thereby change the interpretation of the results. Another alternative hypothesis is that DTEP could be disrupting the completion of autophagy, causing a buildup of LC3 within the nucleus without subsequently increasing autophagic flux, although this is unlikely based on the LC3 turnover experiments. Nuclear preservation may in turn result in an increased potential for oncogenic transformation and the onset of cancer^7^. Future studies utilizing these tools will allow us and others to unravel the complex mechanisms underlying nucleus-associated autophagy.

Now that nucleus-associated autophagies have been generally characterized, there are three important directions in which further research and investigation may continue. First, the identity of the sensors and signaling networks that activate these selective autophagies are still unknown, and substrate targeting is not understood. The egress of nuclear material via autophagy has now been described in more detail^7^, but the mechanism by which the autophagic machinery enters the nucleus is unclear. Previous studies^35^ suggest LC3 enters the nucleus via passive diffusion, but there may be additional pathways identified with the new data that has recently emerged. Previous research states nucleo-cytoplasmic abundance of GFP tagged LC3 remains independently regulated from nuclear export mechanisms. Though LC3 contains a presumed nuclear export signal (NES), it does not necessarily regulate the export of EGFP-LC3 from the nucleus under steady state conditions^35^. In the setting of importin knockdown (IPO8 specifically), we hypothesize some LC3 that might have other otherwise traversed the nuclear envelope was unable to enter the nucleus, implying a partial mode of specific transport by IPO8. Further work will allow us to more carefully explore these mechanisms.

Our second goal queries the role of nucleus-associated autophagy in disease, with a focus on cancer and neurodegenerative diseases such as amyotrophic lateral sclerosis (ALS). Various mutations, including the ALS *C9orf72* mutation, are known to result in nuclear stress due to RAN proteotoxic aggregates and, perhaps more interestingly, the buildup of antisense RNA foci within the nucleus^37^. The implication of nucleophagy as a potential disposal system for these mis-incorporated products may be critical for the survival of cells in the onset of these diseases. *C9orf72* mutants are known to have irregularities in the structure and stability of the nuclear envelope directly related to the length of the repeat, which ultimately leads to degeneration of neuronal tissue and failure of the nuclear pore to allow effective nucleocytoplasmic transport^14^. Given that recent findings suggest the direct connection between *C9orf72* and an increase in autophagic flux^38^, the next step would be to investigate the presence and potential role of nucleophagy in cell survival in the setting of ALS and related diseases.

Autophagy as an active cellular process is highly implicated in the setting of oncogenic stress, as has been previously described. Selective autophagy of the nucleus appears to be particularly relevant in the setting of oncogenic stress, allowing cells to survive multiple adverse factors. In the setting of nuclear stress, selective autophagy is able to mediate and remove damaged DNA, activated oncogenes, and proteotoxic aggregates, therefore providing the cancer cells stability^39^. Previous research has highlighted the role of immune cells (human peripheral blood lymphocytes) in inducing autophagy to promote tumor cell survival and proliferation^34^, and dysfunctional autophagy is known to induce tumorigenesis^40^. Further questions remain regarding the makeup of the tumor microenvironment and the role nucleus-associated autophagies plays in mediating key players including stromal, immune, and endothelial cells, all which may impact autophagic flux. Furthermore, we would like to ask how this cell-mediated behavior can regulate crosstalk within the physiological environment to activate and modulate pathways, respond to metabolic stress, and maintain homeostasis. Ultimately, looking at these heterotypic interactions will give better intuition for therapeutic targets that modulate both autophagy and the tumor microenvironment to hinder cancer progression.

## Acknowledgements

The authors would like to thank Nazila Shafagati for spearheading these experiments and Sara Barlow, Anna Maximova, and Mariel Liebeskind for helping see them through to completion. Abigail Reft provided microscope support. We would like to thank the Michael Lotze lab for some early guidance. Thanks to the University of Miami TEM facility, specifically Ji Lin, for images, Dr. Jeffrey S. Prince, for advising and equipment, Dr. James Baker, for advising and Justin Ma, for assistance. Finally, funding was provided by a collaborative EVMS-WM grant, grant 2R15DK058028 from the National Institutes of Health and 1120513 from the National Science Foundation, the William & Mary Charles Center, Ferguson Funds, HHMI freshman research fellowship and W&M Honors funding.

## Author contributions

All authors took part in writing and reviewing the manuscript. **DPR**: Contributed to the early planning, execution, and analysis of most experiments. He also contributed to development of microscopy techniques and image analysis. **LK**: Contributed to concept, planning, and execution of experiments in Figure 2, Figure 4, and was also influential in the ideas and planning of much of the rest of the paper. **DH**: Contributed to the planning, execution and analysis of all sections. **EC, MJV:** Were both a major part of the experimental design and execution of experiments that led to Figure 1,2,4. **CA**: Contributed to protein analysis of Figures 1 and 3 **JZ**: Planned and executed much of Figure 3. **MV:** Contributed to Figure S3, S5. **LA**: Contributed to the overarching concepts, helped steer the paper, and provide numerous resources. **WJB**: Planned the paper and was part of all the analysis, figure design, and writing.

## Data availability

The datasets generated and analyzed with this study are available from the corresponding author upon reasonable request.

