## Supplemental Figures and Tables for "Modulating Intra-Nuclear LC3 with Small Molecules Rescues Cells from a Docetaxel-Induced Phenotype"

### TITLE PAGE

**Full title**

William J Buchser (Corresponding)

Department of Genetics, Washington University School of Medicine

4515 McKinley Ave

St Louis, MO USA

**Running Title (Head)**

Chemical Nucleophagy Modulators

**Supplemental Figures**

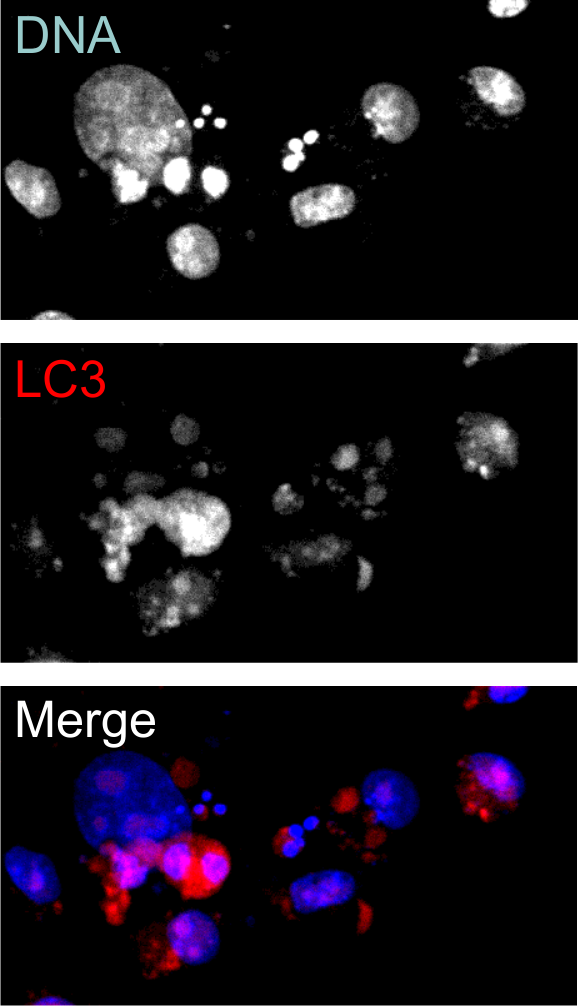

S**upplemental Figure S1. LC3 and Micronuclei.** Micrographs taken with confocal microscopy showing micronuclei with dense LC3 staining. LC3 is frequently perinuclear and in association with micronuclei.

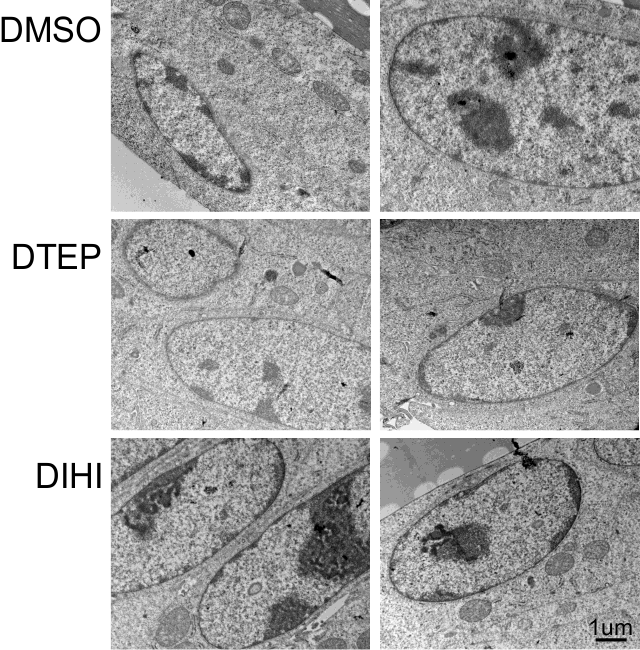

**Supplemental Figure S2. Ultrastructural View of Nucleus with DTEP/DIHI.** Electron micrographs of cells treated with compounds DTEP and DIHI reveal little to no difference in structure from DMSO controls.

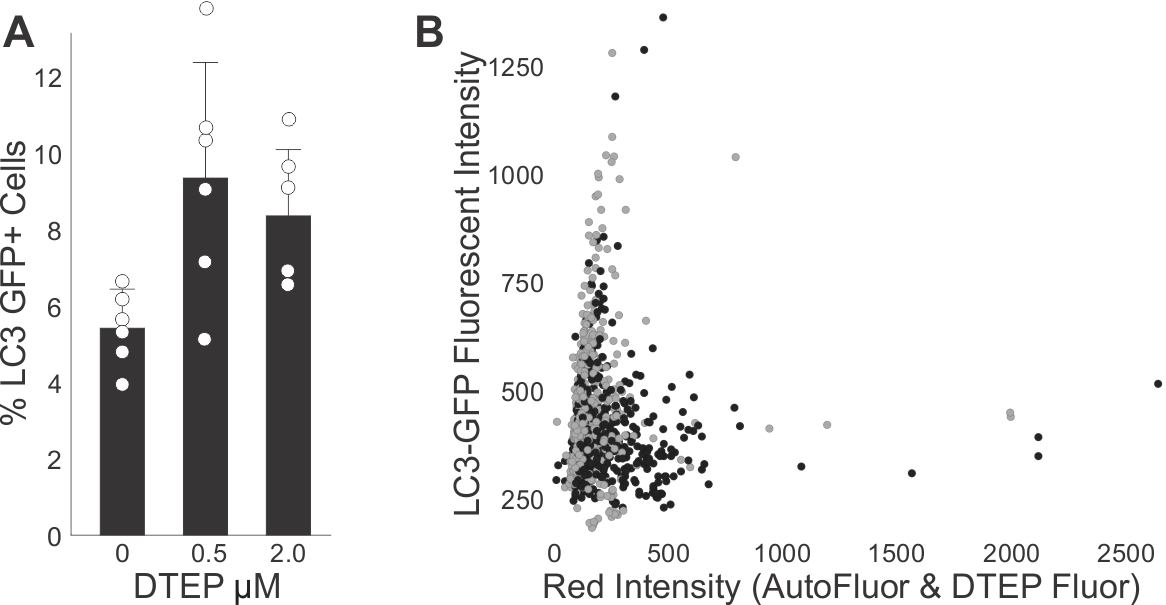

**Supplemental Figure S3. Prevalence of LC3-GFP after the addition of DTEP.** HCT116 cells were infected with LC3-GFP Lentivirus and allowed to grow for 24 hours.  This infection resulted in a physiologically relevant distribution of LC3-GFP (unlike other constructs where the high level of overexpression resulted in a non-physiologic nuclear expression in all the cells). The cells were then imaged and the nuclear LC3 was measured. **A)** The fraction of cells with nuclear LC3-GFP increased after the addition of DTEP (ANOVA p=0.0142, n=6). Individual white markers show the data from an individual image, and the bars show the average for all the images in that condition.  **B)**  Nuclear intensities of each of the cells in the DTEP-treated conditions. There was no correlation (r^2^ = 0.001) between cells that happened to be marked by DTEP itself, and the cells with high nuclear LC3-GFP.  Black markers are cells treated with 2.0 µM DTEP, while gray was 0.5 µM. HCT116 cells were plated for 24 hours then DTEP was added for 1 hour only then the virus (2µl of 1x10^4^ infection units/µl) was added 18 hours before imaging.

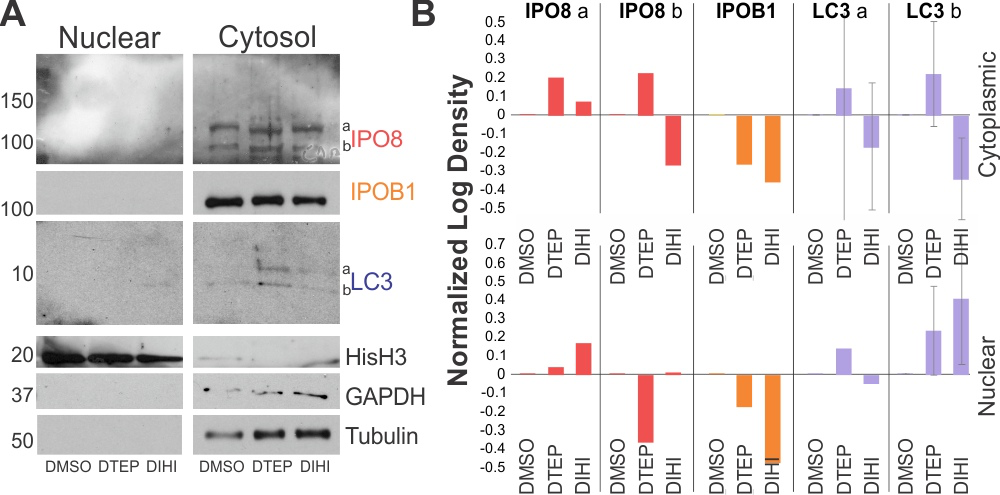

**Supplemental Figure S4. Protein Analysis of Fractions. A)** Western blot analysis of nuclear and cytosolic fractions from renal cells treated for 4 hours with one of the compounds listed. 12 hours after treatment, cells were lysed and processed to analyze subcellular factions (Subcellular Fractionation Kit, Invitrogen 78840). Western membranes are cropped as delineated by the white gaps. There is little significant difference between the treatments and the concentration of both importins and LC3 in the context of DTEP and DIHI. **B)** Quantification of protein levels in both fractions confirms little significant difference in protein levels.

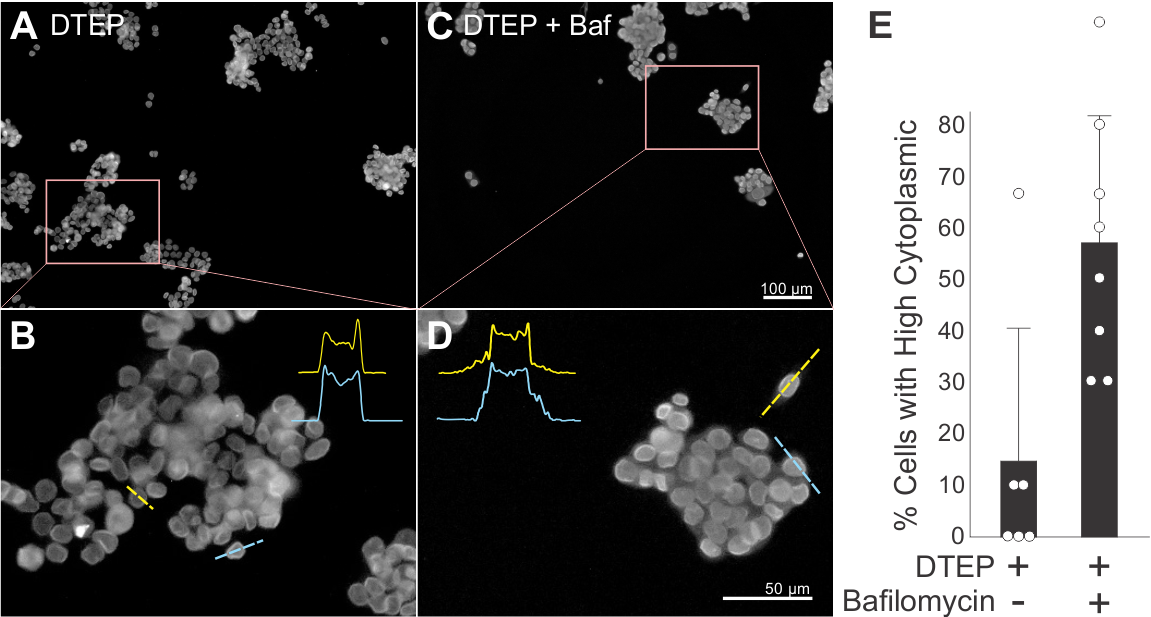

**Supplemental Figure S5. LaminB1 dynamics after DTEP treatment with bafilomycin.** **A, C**) Randomly chosen representative images of laminB1-stained cells that have received 2 µM DTEP +/- 60 nM bafilomycin. **B, D**) Magnified images featuring line tracings of two cells each with corresponding intensity profile plots shown as insets. **E**) Bar chart showing how bafilomycin affects the percentage of cells with cytoplasmic laminB1 (ANOVA p = 0.017). Individual white markers indicate the data from an individual image, and the bars show the average for all the images in that condition. There is a significant increase in the number of cells with cytoplasmic laminB1 when bafilomycin is present. HCT116 Cells were plated, and DTEP was added 24 hours later. After 2 hours, bafilomycin was added then the plates were fixed after an additional four hours. LAMP2 was punctate, and there was a slight increase in co-localization with LaminB1 due to the cytosolic movement of the LaminB1.

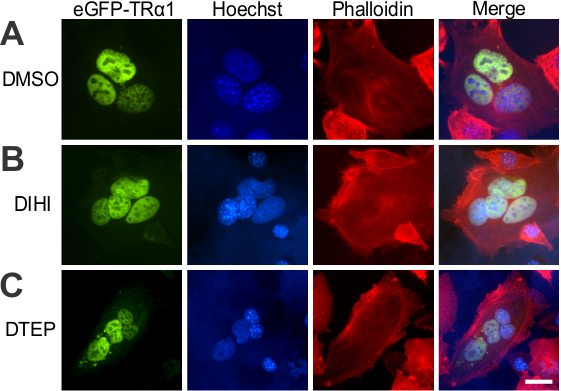

**Supplemental Figure S6. Neither DTEP nor DIHI inhibit nucleocytoplasmic shuttling.** In this heterokaryon assay, nuclei are examined for their ability to transport TRα1 to the nucleus. Cells with no GFP-TRα are merged with cells that express TRα. If shuttling is intact, then all the nuclei in the heterokaryon will contain the green GFP-TRα, as is the case here. All panels are epi-fluorescent micrographs, with the first column the green channel showing GFP-tagged TRα1. The second column is a Hoechst stain for nuclear DNA, marking multiple nuclei within the fused cell. The HeLa nucleus (diffuse blue staining) is distinguished from the NIH/3T3 nucleus (speckled) by differential coloration with Hoechst. The third column shows the extent of the cell with the actin stain phalloidin. The fourth column is a three-color merge. Rows indicate three treatments for 4 hours before the cells were fused. **A**) DMSO control **B**) DIHI **C**) DTEP. All nuclei within the fused cell have been able to transport the receptor, regardless of the treatment. Scale bar 10 µm.

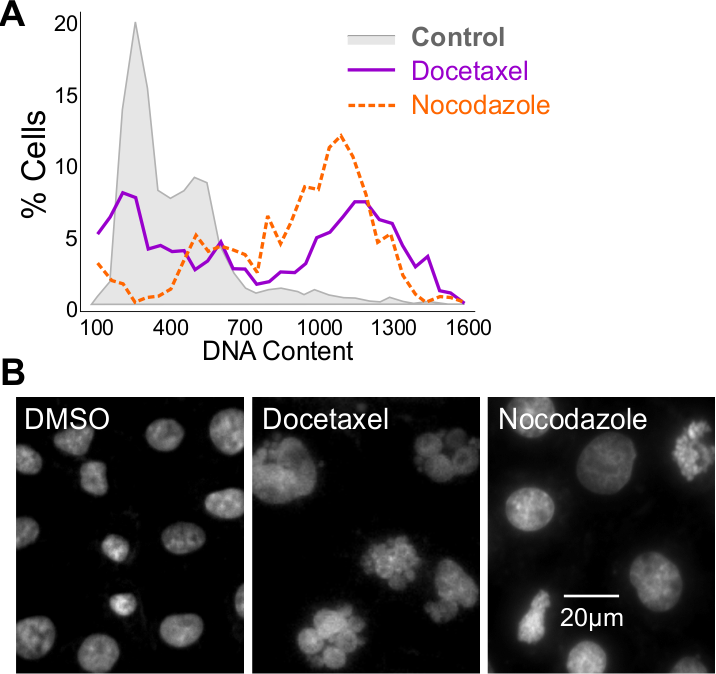

**Supplemental Figure S7.** **Docetaxel-Induced Mitotic Disruption**. **A)** Histograms showing the distribution of individual cells on a scale of integrated blue fluorescence intensity, representative of DNA content. Control cells (gray shaded region) have a primarily unimodal population distribution with most cells in the low DNA content range of 400-700 MFU (mostly diploid nuclei). Nocadozole treated cells (orange dotted line) show a bimodal distribution with more cells in the 800-1400 MFU range, indicating polyploidy. Docetaxel treated cells (purple solid line) appear to show a multimodal distribution of cells with abnormal, non-multiplicative DNA content, likely illustrating mitotic disruption and aneuploidy. **B**) Example micrographs of Hoechst-stained nuclei in DMSO, docetaxel and nocodazole treated 786-0 cells. The middle panel shows the best example of MDIP (mitotic-disruptor induced polyploidy). Note that compared to the DMSO control, docetaxel-treated cells show a distinct lobe-like morphology whereas nocodazole-treated cells are larger than DMSO control nuclei.

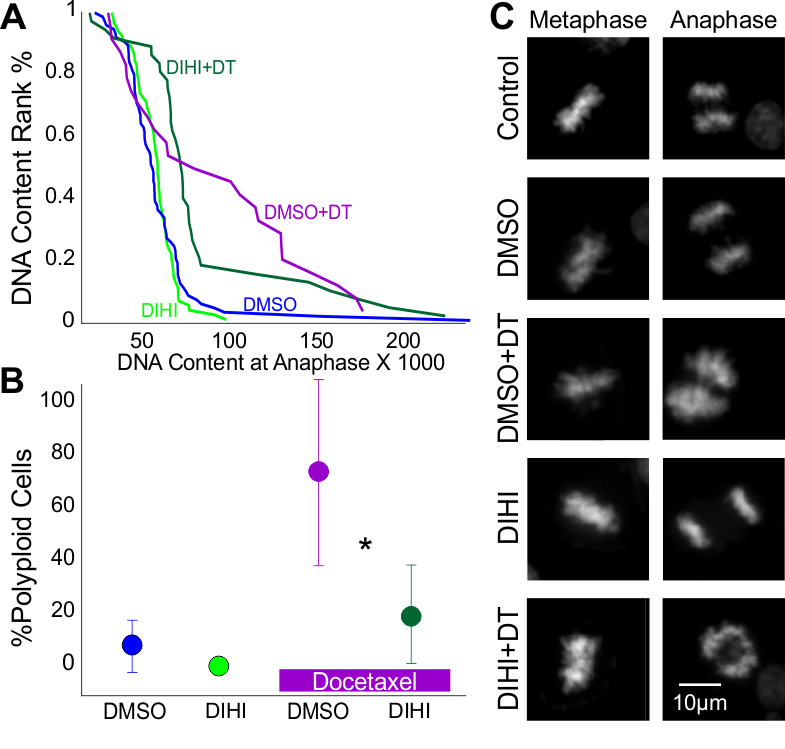

**Supplemental Figure S8. Validating polyploidy with colcemid treatments**

**A)** Cumulative histogram of DNA content (integrated intensity of Hoechst) during anaphase. Individual cells are ranked by their anaphase DNA content. Cells from multiple replicates are pooled. **B)** Effect of DIHI in the presence of Docetaxel on the fraction of polyploid cells (anaphase DNA content > 80 in A). Each point is the average of 5-6 replicates. ANOVA p = 0.007, F=5.802. **C)** Representative images of metaphase and anaphase nuclei (in 786-0 cells) from the various experimental conditions. Cells were halted with 0.1 µg/ml of Colcemid.

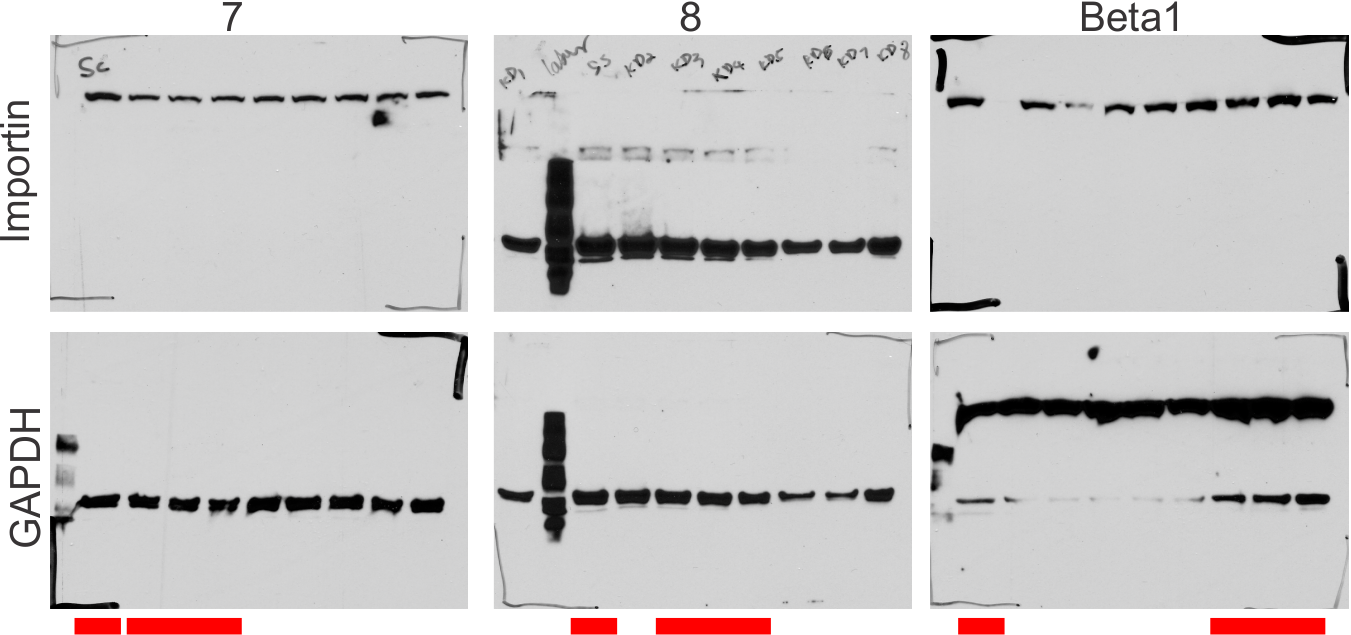

**Supplemental Figure S9. Pre-Cropped Western Blot Membranes from Figure 3.**

First row shows the membranes that were probed with anti-importin antibodies. The bottom row shows the membranes with anti-GAPDH antibodies. The red rectangles below the membranes indicate the regions that were used for the final figure.
